# Synaptic control of DNA-methylation involves activity-dependent degradation of DNMT3a1 in the nucleus

**DOI:** 10.1101/602151

**Authors:** Gonca Bayraktar, PingAn Yuanxiang, Guilherme M Gomes, Aessandro D Confettura, Syed A Raza, Oliver Stork, Shoji Tajima, Ferah Yildirim, Michael R. Kreutz

## Abstract

DNA-methylation is a crucial epigenetic mark for activity-dependent gene expression in neurons. Very little is known how synaptic signals impact promoter methylation in neuronal nuclei. In this study we show that protein levels of the principal *de novo* DNA-methyltransferase in neurons, DNMT3a1, are tightly controlled by activation of N-methyl-D-aspartate receptors (NMDAR) containing the GluN2A subunit. Interestingly, synaptic NMDAR drive degradation of the methyltransferase in a neddylation-dependent manner. Inhibition of neddylation, the conjugation of the small ubiquitin-like protein NEDD8 to lysine residues, interrupts degradation of DNMT3a1 and results in deficits of promoter methylation of activity-dependent genes, synaptic plasticity as well as memory formation. In turn, the underlying molecular pathway is triggered by the induction of synaptic plasticity and in response to object location learning. Collectively the data show that GluN2A containing NMDAR control synapse-to-nucleus signaling that links plasticity-relevant signals to activity-dependent DNA-methylation involved in memory formation.

## Introduction

It is widely believed that rapid and reversible DNA methylation is essential for the stability of long-term memory but still very little is known how synaptic signals can induce changes in DNA-methylation to elicit enduring alterations in plasticity-related gene expression (Bayraktar & Kreutz, 2018a, b; Campbell & Wood, 2019; Day & Sweatt, 2010; Miller & Sweatt, 2007; Guo *et al*, 2011). In addition, aberrant DNA methylation has been implicated in neuropsychiatric diseases including schizophrenia, bipolar, and major depression disorders (Bayraktar & Kreutz, 2018; Mill *et al*, 2008; Murgatroyd *et al*, 2009). NMDAR signaling to the nucleus is instrumental for learning and memory formation and is altered in schizophrenia as well as other neuropsychiatric disorders (Zhou & Sheng, 2013; Paoletti *et al*, 2013). However, a mechanistic link between NMDAR signaling and DNA-methylation is currently missing.

Compelling evidence exists for learning-induced *de novo* DNA methylation with several studies showing the necessity of active DNA methylation as well as demethylation particularly in the hippocampus during memory consolidation (Bayraktar & Kreutz, 2018; Oliveira, 2016; Kaas *et al*, 2013; Rudenko *et al*, 2013). One of the target genes is the brain-derived neurotrophic factor (BDNF), which undergoes promoter-specific DNA demethylation in the CA1 region of the hippocampus during memory consolidation (Lubin *et al*, 2005). The underlying signaling machinery is also here not well understood and it is still essentially unclear how synaptic signals conveyed to the nucleus impact mechanisms of DNA methylation and demethylation of the *Bdnf* promoter. DNMT3a1 is the major *de novo* DNA methyltransferase in brain and plays a documented role in activity-dependent DNA methylation (Feng *et al*, 2005; Feng *et al*, 2010). Along these lines impaired spatial learning and memory and attenuated CA1 long-term potentiation (LTP) have been reported following a forebrain specific DNMT gene knockout in principal neurons (Feng *et al*, 2010; Morris *et al*, 2014).

Finally, it is nowadays widely accepted that memory consolidation as well as synaptic plasticity not only rely on *de novo* protein synthesis but also protein degradation (Jarome & Helmstetter, 2013; Karpova *et al*, 2006). Proteasomal degradation of proteins in neurons has been studied mainly in the context of ubiquitylation and sumoylation, whereas neddylation – the attachment of the small ubiquitin-like peptide neural precursor cell-expressed developmentally down-regulated gene 8 (NEDD8) – has not been much investigated. Here we show that activation of synaptic GluN2A-containing NMDARs drives the neddylation-dependent proteasomal degradation of the principal *de novo* DNA-methyltransferase in the adult brain DNMT3a1. Collectively the data point to a mechanism that allows for the synaptic control of nuclear DNMT3a1 protein levels thereby creating a time window for reduced de novo DNA-methylation at a subset of target genes. This signaling pathway highlights how synapse-to-nucleus signaling might directly impact on DNA-methylation and memory consolidation.

## Results

### Synaptic activity controls levels of DNMT3a1 in neuronal nuclei

DNMT3a1 is the major de novo DNA methyltransferase expressed in the adult brain (Feng *et al*, 2005). When we first addressed the cellular localization of the enzyme using an antibody that recognizes an N-terminal region (Sakai *et al*, 2004) specific for DNMT3a1 (Fig EV1A to D), we found a prominent nuclear localization in hippocampal primary neurons and much fainter staining hardly above background in astrocytes (Fig 1A), indicating that DNMT3a1 is mainly expressed in neurons. This finding prompted us to asked next whether synaptic activity might regulate nuclear DNMT3a1 protein levels. When we induced burst firing of excitatory synapses in hippocampal primary neurons with the GABA-A receptor antagonist bicuculline (bic) and the potassium channel blocker 4-amino-pyridine (4-AP), we observed a prominent reduction in the nuclear immunofluorescence of DNMT3a1 (Fig 1B and C), a finding that was confirmed by quantitative immunoblot analysis of cell lysates from cortical primary neurons (Fig 1D to F). Enhancing excitatory activity for 10 min with bic/4-AP was sufficient to reduce DNMT3a1 immunofluorescence (Fig 1G and H), while excitotoxicity in hippocampal primary neurons elicited with bath application of 100 µM NMDA for 10 minutes – a protocol that results in extrasynaptic NMDAR activation and delayed cell death (Dieterich *et al*, 2008) – had no effect (Fig 1G and H).

**Figure 1.**
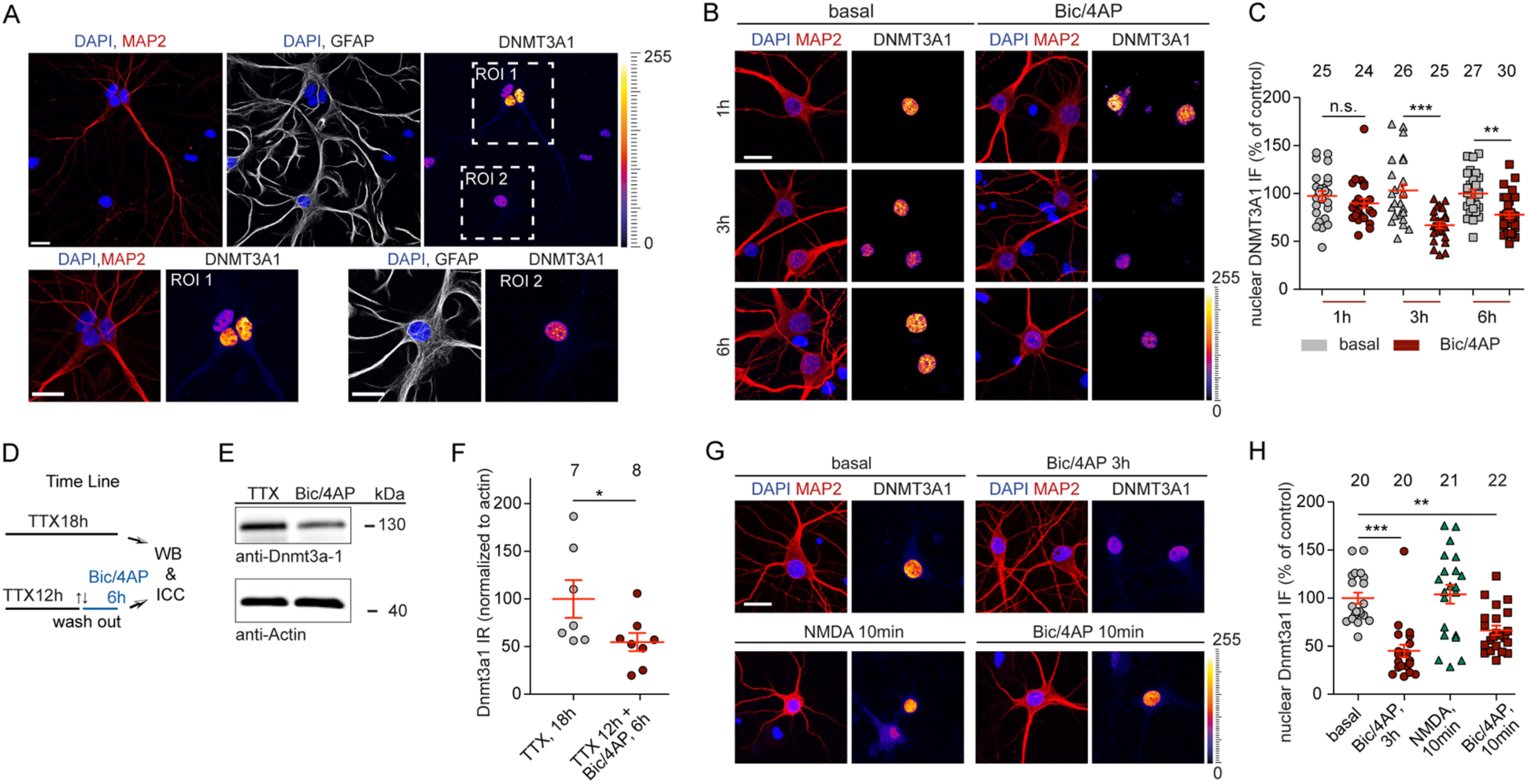
Synaptic activity regulates DNMT3a1 protein levels in neurons. **A** Nuclear DNMT3a1 immunofluorescence is prominent in MAP2-positive neurons but much less in GFAP-positive astrocytes at 15 DIV hippocampal cultures. Scale bars are 20 µm. **B** and **C** Down-regulation of DNMT3a1 protein levels in 14-15 DIV hippocampal primary neurons following treatment with bic/4AP for 1h, 3h or 6h as evidenced by quantitative immunocytochemistry. Scale bar is 20 µm. Students t-test **p<0.01, ***p<0.001. **D** to **F** DNMT3a1 protein levels are decreased following removal of tonic inhibition in DIV 21 cortical neurons. β-Actin was used as an internal control for normalization. Student’s t-test *p<0.05. **G** and **H** Treatment of neurons with bic/4AP for 10 min or 3 h was equally effective to reduce nuclear DNMT3a1 immunofluoresence. Application of 100 µM NMDA for 10 min had no effect. Students t-test **p<0.01, ***p<0.001.

### Synaptic GluN2A containing NMDAR drive degradation of nuclear DNMT3a1

The activity-dependent degradation of DNMT3a1 was blocked in the presence of the competitive NMDAR antagonist 2-amino-5-phosphonopentanoic acid (APV) (Fig. 2A and B). Interestingly, the application of the antagonist NVP-AAM077 at low doses, that mainly target di-heteromeric GluN2A-containing NMDARs (Auberson *et al*, 2002), completely prevented degradation (Fig. 2C and D). In addition, shRNA-induced protein knockdown (Fig EV2A and B) of GluN2A confirmed a specific requirement for NMDAR containing this subunit to elicit degradation of DNMT3a1 (Fig 2E and F), whereas application of the GluN2B antagonist ifenprodil had no effect (Fig 2G and H). Co-application of the L-type Ca^2+^-channel blocker nifedipine (Fig EV2C and D) or the CaMK inhibitor KN93 to the stimulation buffer also hindered nuclear DNMT3a1 degradation (Fig EV2E and F). Taken together, these experiments indicate that brief activation of synaptic GluN2A containing NMDAR, probably by induction of dendritic action potentials and nuclear calcium waves, is a potent stimulus to control DNMT3a1 protein levels in the nucleus.

**Figure 2.**
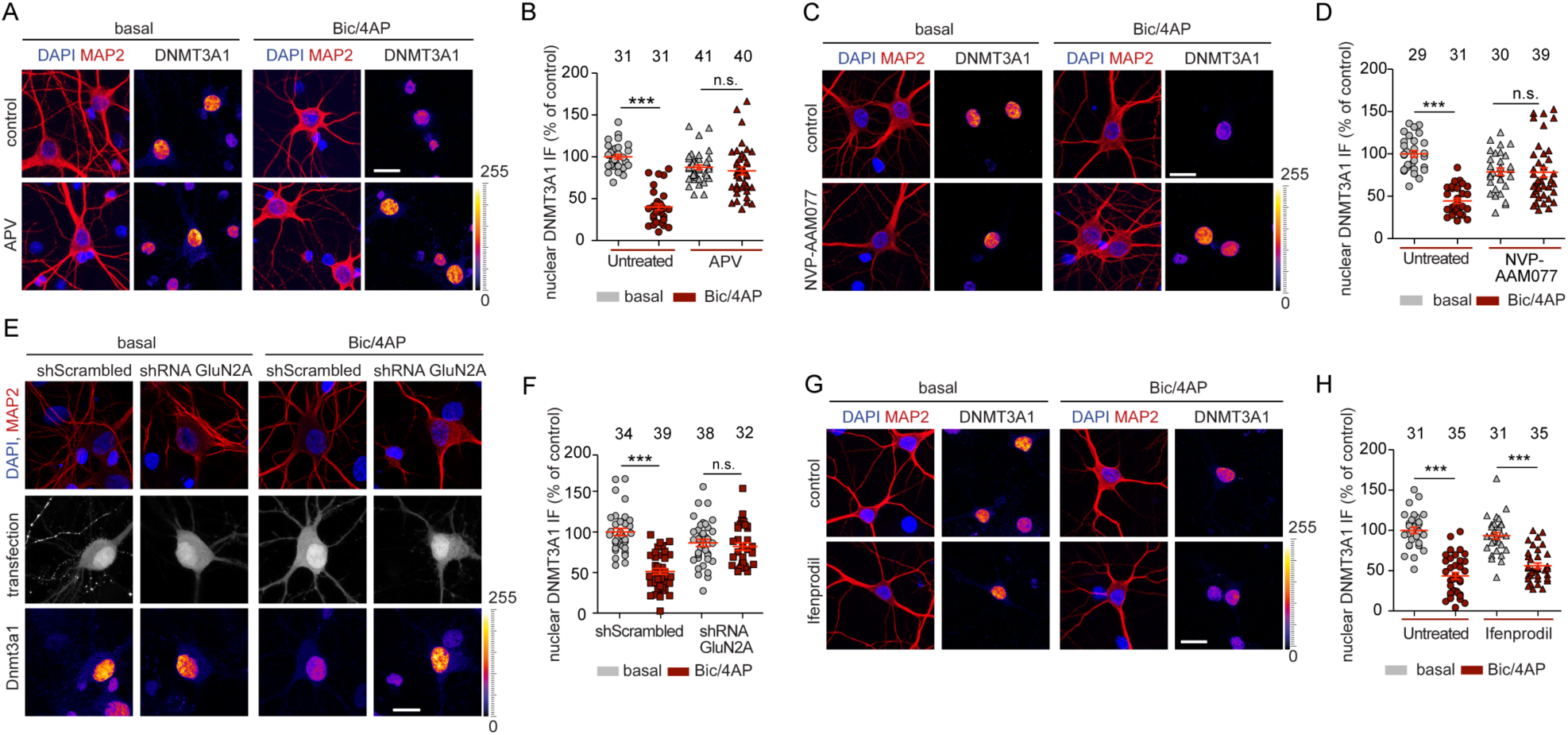
GluN2A-dependent NMDAR signaling regulates nuclear DNMT3a1 protein levels in 14-15 DIV hippocampal primary neurons. **A** and **B** Treatment with the NMDAR antagonist APV (20 µM) prevented the reduction in the nuclear levels of DNMT3a1 following synaptic stimulation by bic/4AP for 6 h. **C** and **D** Treatment with the GluN2A inhibitor NVP-AAM077 (50 nM) prevented the reduction in the nuclear levels of DNMT3a1 following synaptic stimulation by bic/4AP for 6 h. **E** and **F** shRNA-based knockdown of GluN2A in the presence of synaptic stimulation with bic/4AP for 6 h prevented the DNMT3a1 degradation following synaptic stimulation by bic/4AP for 6 h. **G** and **H** Treatment with the GluN2B subunit inhibitor ifenprodil (10 µM) in the presence of synaptic stimulation with bic/4AP for 6 h did not prevent the reduction in the nuclear levels of Dnmt3a1. Scale bars, 20 µm. Two-way ANOVA followed by Bonferroni’s post hoc test. ***P<0.001, n.s. not significant.

### Proteasomal degradation of DNMT3a1 requires neddylation

We next examined which mechanisms might contribute to DNMT3a1 down-regulation and found that the proteasome inhibitors MG132, Lactacystin or Carfilzomib, which all operate via different mechanisms (Fenteany *et al*, 1995; Lee & Goldberg, 1998; Meng *et al*, 1999), completely abolished the effect of GluN2A stimulation (Fig EV3A to F). Since we found no concomitant alteration of DNMT3a1 mRNA levels (Fig EV3G), these data suggest that proteasomal degradation controls the protein levels of the enzyme in an activity-dependent manner.

Potential mediators of DNMT3a1 degradation are members of the family of Cullin proteins (Sarikas *et al*, 2011). Cullin family members combine with RING proteins to form Cullin-RING E3 ubiquitin ligases (Petroski & Deshaies, 2005) and neddylation is a prerequisite for their activation in the nucleus. Neddylation has been studied only recently in neurons (Vogl et al., 2015; Scudder & Patrick, 2015) and the subcellular distribution of NEDD8 in neurons has not been determined yet. We found that NEDD8 is abundantly localized in the nucleus of hippocampal primary neurons, whereas immunofluorescence intensity was much weaker at synapses (Fig 3A). We observed heterologous co-immunoprecipitation of DNMT3a1 with CUL4B, CUL1, CUL3, CUL4, CUL7, but not with CUL2 and CUL5 from HEK293T cell lysates (Fig EV4A). In addition, we found that poly-ubiquitination of DNMT3a1 was elevated following forced expression of CUL4B in HEK293T cells (Fig EV4B to D), which was chosen as an example because of its high expression in brain. Conversely shRNA-based CUL4B protein knockdown resulted in reduced DNMT3a1 poly-ubiquitination (Fig EV4E). The neddylation inhibitor MLN4924 selectively inhibits the NEDD8-Activating-Enzyme (NAE) at very low concentrations (Soucy *et al*, 2009). Accordingly, poly-ubiquitination of immunoprecipitated DNMT3a1 was reduced in cells that were subjected to MLN4924 treatment (Fig EV4F). Moreover, neddylated CUL4B was co-immunoprecipitated with DNMT3a1 (Fig EV4G).

**Figure 3.**
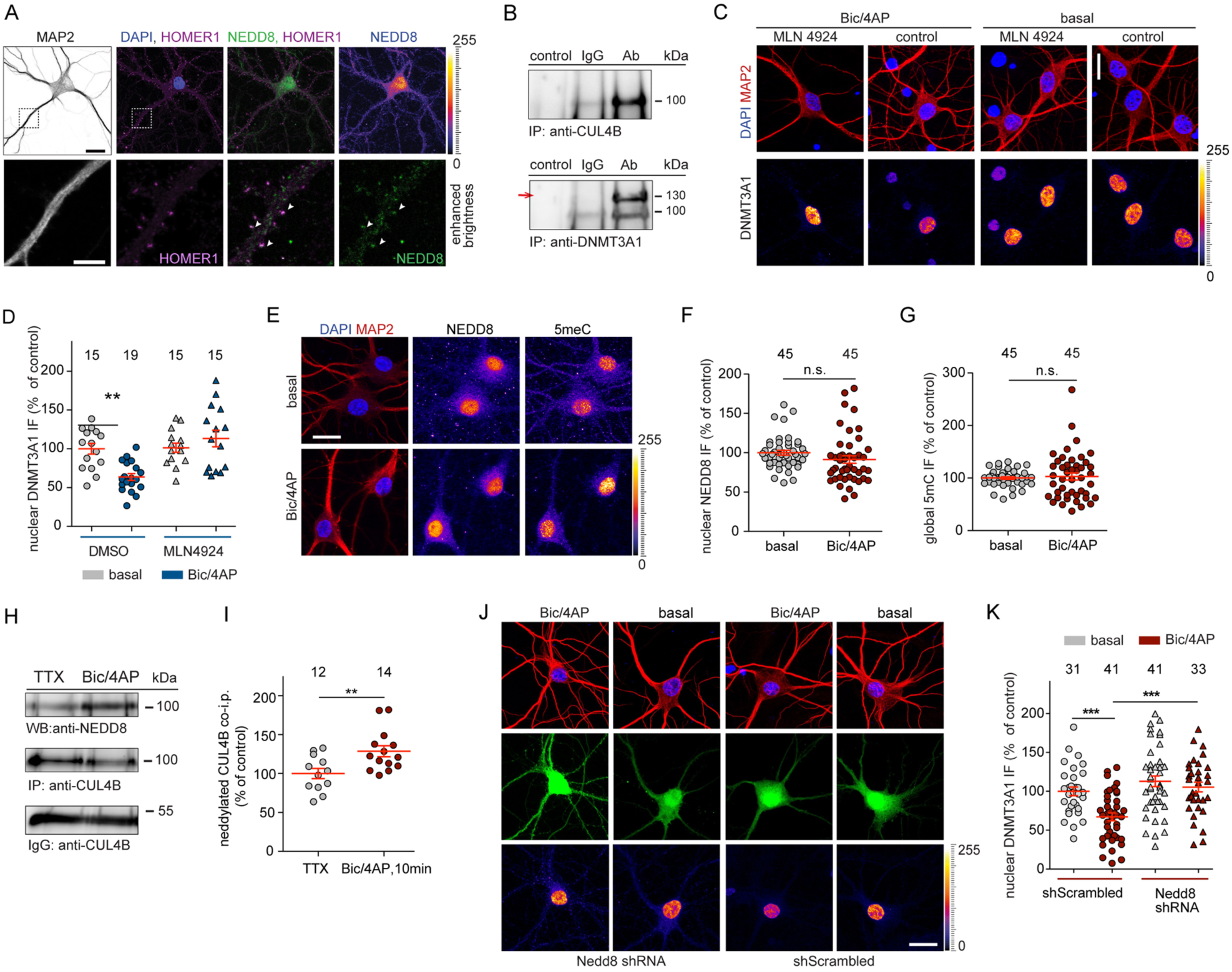
DNMT3a1 degradation is a neddylation dependent process. **A** NEDD8 is abundantly expressed in the nucleus of 16 DIV hippocampal neurons whereas weaker staining was observed at synapses. Scale bar 20 µm (top panel), 5 µm (lower panel). **B** Immunoprecipitation of CUL4B protein from nuclear extracts of cortical primary neurons (top panel) results in co-precipitation of Dnmt3a1 (lower panel, indicated with an arrow). **C** and **D** Hippocampal primary neurons were treated with bic/4AP for 6 h in the presence or absence of MLN4924 (5 nM). Scale bars, 20 µm. Quantitative immunocytochemistry revealed that blocking neddylation prevents Dnmt3a1 degradation. **E** Representative immunofluorescence images of nuclear NEDD8 and total cytosine methylation (5mC) at basal conditions or after treatment with bic/4AP for 6 h. Scale bar is 20 µm. **F** and **G** Upon synaptic activation nuclear NEDD8 and 5mC levels remained unchanged. Student’s t-test. n.s. not significant. **H** and **I** CUL4B was immunoprecipitated from nuclear extracts of primary cortical neurons and neddylated CUL4B was quantified. Following 10 min of synaptic stimulation the amount of neddylated CUL4B was increased. Student’s t-test **p<0.01. **J** and **K** shRNA-based knockdown of NEDD8 in hippocampal primary neurons occluded the DNMT3A1 degradation following 10 min-long bic/4AP treatment and fixation of cells 3 h after washout of the drug-containing-media. Two-way ANOVA followed by Bonferroni’s post-hoc test. ***p<0.001, scale bar, 20 µm.

We next asked whether neddylation of Cullins might be involved in controlling the activity-dependent DNMT3a1 proteasomal degradation in neurons. Co-immunoprecipitation experiments of endogenous proteins extracted from cortical primary neurons using pars pro toto a CUL4B specific antibody revealed that neuronal CUL4B and DNMT3a1 might be in one complex in vivo (Fig 3B). Acute treatment of hippocampal primary neurons with concentrations of MLN4924 as low as 5 nM prevented DNMT3a1 degradation (Fig 3C and D). Of note, acute treatment of primary neurons for six hours with either a low (5 nM) or even a high dose of MLN4924 (1 µM) did not alter the total number of spines (Fig EV5A and B) as it was reported previously with the higher concentration and long-term treatment (Vogl et al., 2015; Scudder & Patrick, 2015). Moreover, we observed that nuclear NEDD8 staining intensity was not altered following bic/4AP treatment (Fig 3E and F), which excludes the possibility that activity-dependent nucleocytoplasmic shuttling of NEDD8 contributes to DNMT3a1 degradation. In addition, 5meC immunocytochemistry revealed that no gross quantitative alterations in global DNA methylation occur following sustained stimulation of synaptic GluN2A containing NMDARs (Fig 3E and G). However, enhanced synaptic activity resulted in rapidly increased neddylation of CUL4B (Fig 3H and I), whereas shRNA knockdown of NEDD8 (Fig EV5C to E) prevented activity-dependent degradation of DNMT3a1 (Fig 3J and K). Collectively, the biochemical, pharmacological and shRNA knock-down experiments show that synaptic activation of GluN2A-containing NMDAR will drive neddylation of Cullin-ligases in the nucleus, which is a prerequisite for DNMT3a1 degradation.

### Synaptic plasticity inducing stimuli elicit DNMT3a1 degradation in a GluN2A-dependent manner

We next addressed whether induction of NMDAR-dependent long-term potentiation (LTP), a form of plasticity that is considered to be a cellular model of learning and memory, involves synaptic control of nuclear DNMT3a1 protein levels. When we induced LTP in the hippocampus with high-frequency stimulation of Schaffer-collaterals (Fig 4A), we found a significant down-regulation of DNMT3a1 protein levels within six hours in the potentiated CA1 region following tetanization of the slices (Fig 4B and C). Moreover, the expression of late-LTP is neddylation-sensitive and field excitatory postsynaptic potentiation slope values returned to baseline within three hours when the slices were treated with the NEDD8 inhibitor MLN4924 (Fig 4D to F).

**Figure 4.**
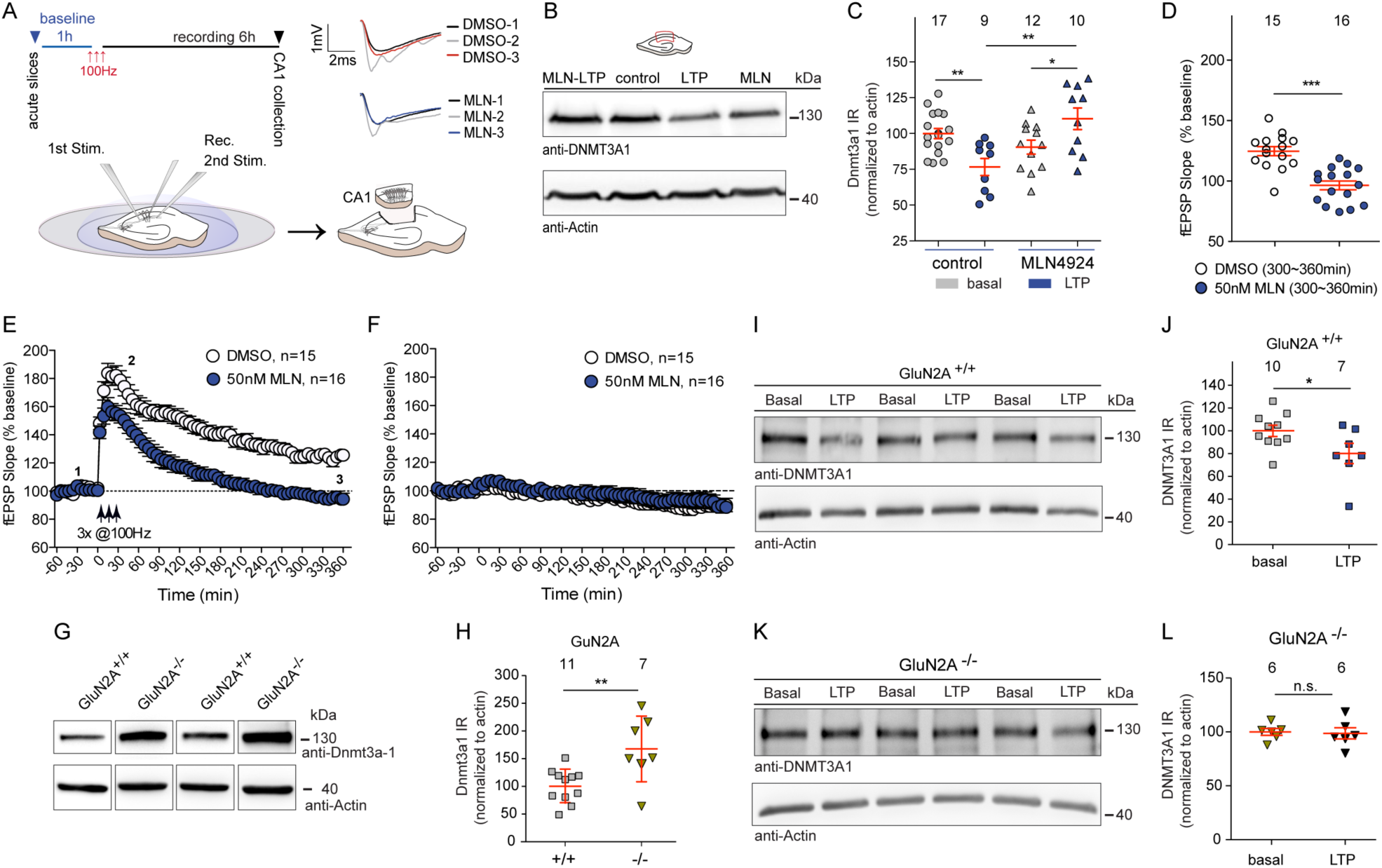
The NEDD8 inhibitor MLN4924 impairs LTP in rat hippocampal slices and degradation of DNMT3a1 upon LTP induction is absent in GluN2A Knock out mice. **A** Schematic presentation of LTP induction by high-frequency stimulation in acute CA1 hippocampal slices. Only the potentiated CA1 region was dissected from individual hippocampal slices 6 h following LTP recordings. Representative traces from recordings at the time points indicated under control and treatment conditions. **B** and **C** DNMT3a1 levels are reduced 6 h after LTP induction and neddylation inhibition prevented the degradation of Dnmt3a1. β-Actin was used as an internal control for normalization. Two-way ANOVA followed by Bonferroni’s post hoc test. **p<0.01, *p<0.05. **D** Averaged fEPSP slopes of the last hour following LTP induction showed significantly reduced LTP in MLN4924 (50nM) treated slices in comparison to slices treated by DMSO. Mann Whitney U-Test ***P<0.001. (**E**) MLN4924 induces significantly impaired LTP as compared to controls. *p<0.05 and **P<0.01. (**F**) Baseline recordings revealed no alterations. **G** and **H** GluN2A knockout (KO) mice show higher DNMT3a1 protein levels in the CA1 region of hippocampus compared to the age-matched wild type (WT) control mice. Students t-test **p<0.01. **I** to **L** Quantitative immunoblotting either from the CA1 of WT (**I** and **J**) or GluN2A KO (**K** and **L**) mice revealed no reduction in the GluN2A KO mice following 3 h of LTP recordings unlike reduction seen in the WT control mice tissue. Unpaired student’s t-test. *P<0.05, n.s. not significant.

We next asked whether following the induction of LTP signaling of GluN2A-containing NMDAR to the nucleus is crucial in neddylation-dependent degradation of DNMT3a1. Interestingly, DNMT3a1 protein levels were already clearly higher in hippocampal tissue homogenates of GluN2A knockout mice as compared to wild-type controls (Fig 4G and H). GluN2A knockout mice show reportedly impaired hippocampal LTP (Sakimura *et al*, 1995) but a stronger tetanic stimulation restores the impairment and the saturation level of LTP is unaltered (Kiyama *et al*, 1998). We could replicate these published findings (Fig EV6A to C) and found that despite the induction of LTP with a stronger protocol no reduction in nuclear DNMT3a1 protein levels like in wild-type mice was detectable (Fig 4I to L). Of note, while we observed a negative correlation between the strength of LTP and the magnitude of DNMT3a1 degradation in wild-type control animals (Fig EV6D) no correlation was seen in GluN2A −/− mice despite recovered LTP. Thus, GluN2A signaling in synaptic plasticity and not the induction of LTP as such is instrumental in controlling nuclear protein levels of DNMT3a1.

### Degradation of DNMT3a1 facilitates Bdnf gene expression

In the adult brain brain-derived neurotrophic factor (BDNF) has principal functions in synaptic plasticity, learning and memory (Karpova, 2014). Expression of the *Bdnf* gene is controlled by eight promoters (Aid *et al*, 2007) and among those, particularly promoter IV activity is strongly stimulated by the calcium influx through synaptic NMDARs (Lubin *et al*, 2005; Zheng *et al*, 2011). DNA methylation of the *Bdnf* IV promoter has been studied previously also in the context of neuropsychiatric disorders (Kundakovic *et al*, 2015; Maynard *et al*, 2016). We therefore chose *Bdnf* IV gene expression to test whether activity-dependent degradation of DNMT3a1 might impact DNA methylation of promoters of plasticity-related genes and corresponding gene expression. Quantitative real-time PCR experiments first revealed that Bdnf mRNA expression is increased by enhanced synaptic activity in primary hippocampal neurons and this increase in transcript levels was significantly lower in the presence of MLN4924 (Fig 5A). Comparable results were obtained in acute hippocampal slices following high-frequency stimulation of Schaffer-collaterals (Fig 5B) and importantly, the LTP-induced increase in Bdnf IV transcript levels was reduced in the presence of the NEDD8-inhibitor (Fig 5B).

**Figure 5.**
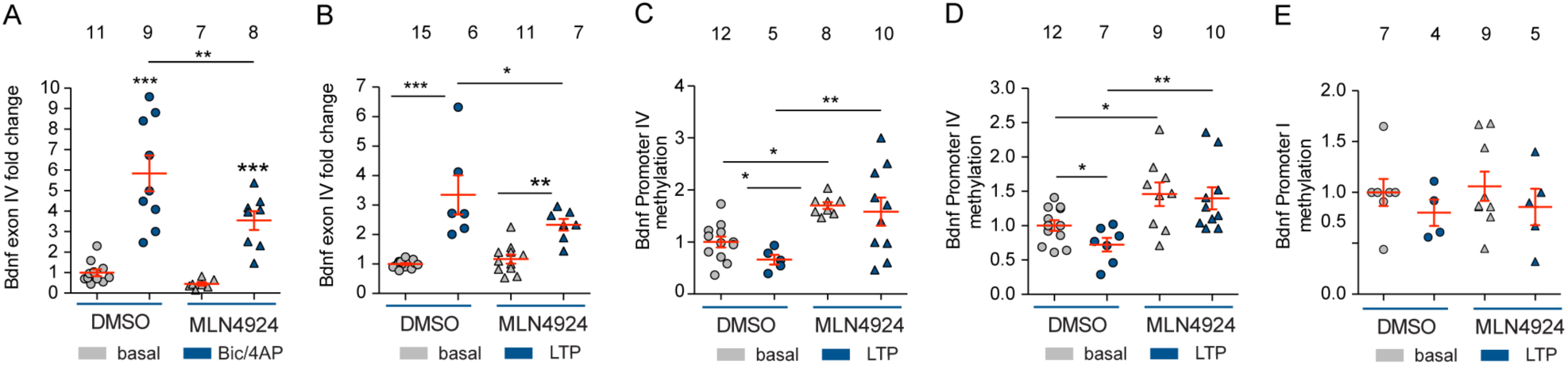
Bdnf IV gene expression and promoter methylation is regulated by neddylation in neurons upon synaptic stimulation. **A** Bdnf exon IV expression was induced in cultured hippocampal primary neurons following bic/4AP treatment for 6h and this induction was lowered by 5 nM MLN4924. Two-way ANOVA followed by Bonferroni’s post hoc test, **P<0.01 ***P<0.001. **B** Bdnf exon IV expression was induced following LTP induction for 6h in CA1 region of Hippocampus and this induction was lowered by 50 nM MLN4924. Two-way ANOVA followed by Bonferroni’s post hoc test, *p<0.05 **P<0.01 ***P<0.001. **C** Methylation analysis based on the restriction of the methylation-sensitive sites showed demethylation of Bdnf promoter IV following LTP induction in CA1 (one-tailed Student’s t-test, t15=1.939, p=0.0358) and promoter methylation was increased when treated with MLN4924 irrespective of LTP induction or basal conditions. Two-way ANOVA followed by Bonferroni’s post hoc test, *p<0.05 **P<0.01. **D** MeDIP-qPCR analysis revealed demethylation of Bdnf promoter IV following LTP induction in CA1 (one-tailed Student’s t-test, t17=2,198 p=0.0421) and promoter methylation was increased when treated with MLN4924 irrespective of LTP induction or basal conditions. Two-way ANOVA followed by Bonferroni’s post hoc test, *p<0.05 **P<0.01. **E** MeDIP-qPCR analysis did not show any alteration in Bdnf promoter I methylation. Two-way ANOVA followed by Bonferroni’s post hoc test.

We next addressed whether increased Bdnf IV mRNA production was correlated with the demethylation of the Bdnf IV promoter in acute slices following LTP induction, as predicted by the degradation of DNMT3a1. First, methylation specific restriction enzyme analysis was performed using primers that span the Bdnf IV promoter sequence possessing three different restriction sites (Fig EV7A). Tetanized CA1 samples revealed a reduction in Bdnf IV promoter methylation, whereas increased promoter methylation was observed for the group that received high-frequency stimulation while being treated with MLN4924 (Fig 5C). A subsequent series of MeDIP-qPCR experiments were performed using tetanized CA1 tissue samples following the induction of LTP and confirmed the change in promoter methylation. Less amplicons were generated with primers targeting the Bdnf IV promoter that cover multiple cytosine residues (Fig EV7B) known to regulate mRNA expression (Fig 5D). More amplicons were detected in MLN4924-treated slices following the induction of LTP (Fig 5D), indicating increased promoter methylation. Among the different Bdnf promoters that were investigated, activity-dependent DNA methylation is particularly prominent for promoter IV, whereas promoter I methylation was not altered following either LTP induction or NEDD8 inhibition with MLN4924 (Fig 5E).

### DNMT3a1 is degraded in the hippocampus as a result of learning

In the final set of experiments, we investigated whether DNMT3a1 degradation occurs *in vivo* as a result of CA1-dependent learning and whether this degradation and memory formation is neddylation-sensitive. Formation of a memory for the spatial location of objects in an open field (Fig 6A) requires synaptic activity of CA1 neurons (Assini *et al*, 2009; Haettig *et al*, 2013) and is responsive to changes in the expression of BDNF (Intlekofer *et al*, 2013; Wang *et al*, 2017). We observed that DNMT3a1 protein levels were reduced in mice three hours following training (Fig 6B and C). DNMT3a1 protein levels returned to control values within six hours, which might reflect less intense synaptic activity as compared to tetanization of slices by high-frequency stimulation (Fig 6D and E). Bilateral intra-hippocampal infusion of MLN4924 into CA1 immediately after object location learning (Fig 6F) resulted in a disturbance of object location memory, as indicated by profoundly reduced discrimination of novel and familiar object locations when compared to mice that received vehicle infusion (Fig 6G). Interestingly, the learning impairment in MLN4924 treated mice was associated with the prevention of DNMT3a1 degradation (Fig 6H and I). Protein levels were significantly higher in mice injected with MLN4924 three hours after training compared to vehicle-injected mice (Fig 6H and I). Six hours following training DNMT3a1 protein levels were no longer different between treatment groups and returned to baseline levels (Fig 6J and K).

**Figure 6.**
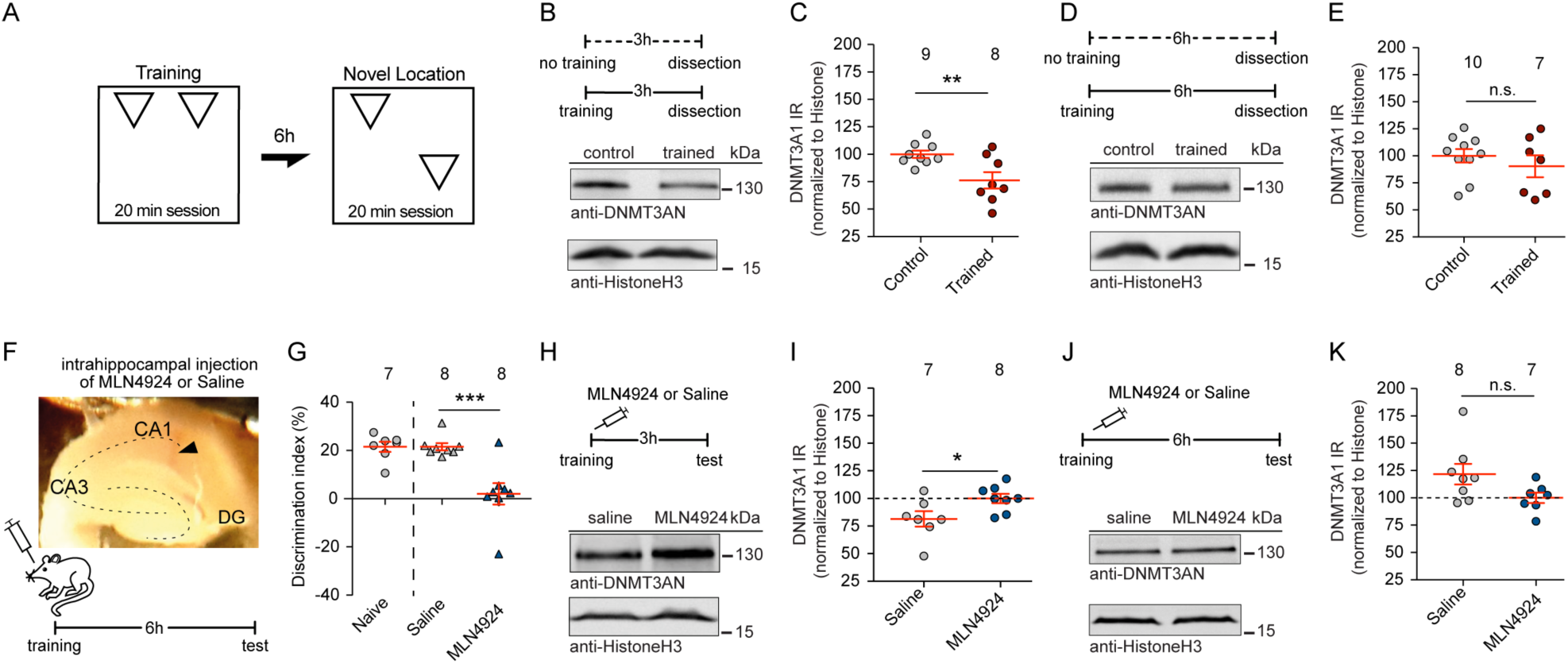
Inhibition of neddylation in CA1 impairs object location memory and the learning-induced degradation of DNMT3a1. **A** Schematic representation of the object location memory experimental protocol. **B** and **C** DNMT3a1 protein levels are reduced 3 h after training. Student’s t-test *p<0.05. **D** and **E** DNMT3a1 protein levels are not changed in trained mice 6 h after training. Histone was used as a loading control for normalization. Student’s t-test n.s. not significant. **F** Injection timeline and representative image of injection site in CA1. **G** Intra-hippocampal infusion of MLN4924 reduces the discrimination index of animals tested in the object location memory test. Unpaired Student’s t-test with Welch correction ***P<0.001. **H** and **I** Learning induced reduction of DNMT3a1 protein levels in trained, but not MLN4924 injected mice 3h after training. Unpaired two-tailed Student’s t-test *p<0.01. **J** and **K** DNMT3a1 protein levels were not altered in trained MLN4924 injected or control mice 6h after training. Histone was used as a loading control for normalization. Data are represented as mean ± S.E.M. Student’s t-test *p<0.05.

## Discussion

Compelling evidence exists for the necessity of active DNA methylation as well as demethylation in the hippocampus during memory consolidation (Bayraktar & Kreutz, 2018a, b; Oliveira, 2016; Kaas et al, 2013; Rudenko *et al*, 2013). However, the underlying signaling machinery is not understood and it is essentially unclear how synaptic signals conveyed to the nucleus impact DNA methylation and demethylation. Here, we show that activation of synaptic GluN2A-containing NMDARs drives the neddylation-dependent proteasomal degradation of the principal *de novo* DNA-methyltransferase in the adult brain DNMT3a1.

The finding that activation of GluN2A containing NMDARs to the nucleus evokes degradation of DNMT3a1 raises several interesting questions about the underlying mechanism of long-distance signaling and the rationale behind it. GluN2A containing NMDAR are in contrast to those containing GluN2B preferentially found at synaptic sites (Wyllie & Hardingham, 2013) and it is possible if not likely that steep and fast synaptic Ca^2+^-influx through these receptors is necessary to elicit nuclear Ca^2+^-responses that in turn enhance neddylation of Cullins. Neddylation as such has not been investigated in any detail in neurons yet and not much information is currently available on how NAE activity itself is regulated. The present study therefore provides first evidence that an NMDAR-derived synaptic calcium signal is coupled to neddylation of Cullins in the nucleus. Two previous reports have shown that blocking neddylation for extended periods of time (in contrast to the administration regime in the present study) leads to reductions in spine size and impairment of synapse maturation in neurons (Vogl *et al*, 2015, Scudder & Patrick, 2015). In addition, neddylation alters synapse function and morphology by directly modifying one of the major synaptic scaffolding proteins PSD95 (Vogl *et al*, 2015). We found that NEDD8 is most abundant in neuronal nuclei and it is tempting to speculate that activity-dependent neddylation might reduce the protein levels not only of DNMT3a1 but also of other nuclear epigenetic modifiers which contribute to object location memory. Along these lines the contribution of neddylation to object location memory seems to be substantial, considering the near-complete removal of the discrimination by the inhibition of neddylation. This is, however, not reflected in the extent of reduction in the levels of DNMT3a1 induced by the behavioral training. Moreover, neddylation is like other posttranslational modifications reversible, which adds potentially another level of regulation for degradation of this methyltransferase. In addition, different efficiencies of proteasomal degradation in neuronal sub-compartments, the necessity for the integration of signaling pathways in the nucleus as well as the complex formation and the potential nuclear export of ubiquitinated DNMT3a1 might account for the relatively slow decline.

Moreover, in this study we propose a long-distance signaling pathway that can provide a potential link between different observations. Converging evidence suggests that NMDAR function in the dorsal CA1 area is critical for novel object location memory (Assini *et al*, 2009; Haettig *et al*, 2013) and increased BDNF expression in the hippocampal CA1 region supports object location learning (Intlekofer *et al*, 2013; Wang *et al*, 2017). We have, therefore, chosen BDNF as a paradigmatic example for our studies, which is one of the target genes that undergoes promoter-specific DNA demethylation in the CA1 region of the hippocampus during memory consolidation (Lubin *et al*, 2005) and impaired spatial learning and memory as well as attenuated CA1-LTP have been reported following a forebrain specific DNMT1 and -3 gene knockout in principal neurons (Feng *et al*, 2010, Morris *et al*, 2014). Aberrant DNA methylation has been implicated in a plethora of studies in neuropsychiatric diseases including schizophrenia, bipolar, and major depression disorders (Bayraktar & Kreutz, 2018; Mill *et al*, 2008; Murgatroyd *et al*, 2009). One of the hallmarks of schizophrenia is a down-regulation of BDNF expression that is associated with the enrichment of 5-methylcytosine at gene regulatory domains within the *Bdnf* promoter (Zheleznyakova *et al*, 2016). Moreover, elevated hippocampal DNMT3a expression has been reported in the postmortem brain of schizophrenia patients (Zhubi et al, 2016).

Collectively our data point to a mechanism that allows for the synaptic control of DNMT3a1 levels and thereby creates a time window for reduced *de novo* DNA-methylation at a subset of target genes. DNMT3a1-mediated methylation has been largely associated with silencing of promoters, which would in turn attenuate activity-dependent gene expression. A shorter splice isoform, DNMT3a2, was shown to associate with transcriptional facilitation of the expression of plasticity-relevant genes presumably via methylation of CpG islands in their promoter and coding regions (Oliveira et al, 2012, 2016). *Dnmt3a2* is an immediate early gene that is identical to *Dnmt3a1* except that it lacks the sequence encoding the N-terminal 219 amino acids of the enzyme, which encompasses the epitope of the antibody that was used in the current study. An intriguing question is whether the increased expression of DNMT3a2 mRNA and down-regulation of DNMT3a1 is induced by the same stimulus, i.e. activation of synaptic GluN2A NMDAR. It is currently unknown whether DNMT3a2 mRNA will be immediately translated, as expected for an immediate early gene. In this case it might not replace DNMT3a1 but independently further facilitate activity-dependent gene expression.

## Materials and Methods

### Primary neuronal culture and drug treatments

Rat cortices and hippocampi were dissected from embryonic day 18 rats (Sprague Dawley). Cells were plated in a density of 30.000 cells per 18 mm coverslip, grown in 1 ml of neurobasal medium (NB, Gibco) supplemented with B27 medium. Primary neurons were kept in Neurobasal medium (NB/GIBCO/Life Technologies) supplemented with B27 (GIBCO/Life Technologies), L-Glutamine (GIBCO/Life Technologies) and penicillin/streptomycin (PAA Laboratories, Pasching, Austria). On day 4 after plating cortical neurons were treated with 5 µM Cytosine D-Arabino-Furanoside (Sigma-Aldrich) in order to prevent proliferation of non-neuronal cells.

Hippocampal neurons were treated at DIV 14-15 with the following drugs for 10 min, 1h, 3h or 6h as indicated in the Results secton: Bicuculline methiodide (50 µM, Tocris, Bristol, UK), 4-Aminopyridine (4AP, 2.5 mM, Sigma-Aldrich, St. Louis, MO), MG132 (30 µM, AG Scientific), Lactacystin (15 µM, Sigma-Aldrich, St. Louis, MO), Carfilzomib (100 nM, UBPBio), NVP-AAM007 (50 nM, Novartis), APV (20 µM, Calbiochem), Ifenprodil (10 µM, Tocris, Bristol, UK), Nifedipine (10 µM, Santa Cruz), KN93 (5 µM, Tocris, Bristol, UK) and MLN4924 (5 nM, Cayman Chemicals, Hamburg, Germany). Hippocampal neurons were treated for 3, 5 or 10 minutes with 100 µM N-Methyl-D-aspartic acid (NMDA, Sigma-Aldrich, St. Louis, MO). Cortical neurons were seeded in T-75 flasks (Thermo Scientific, Rockford, USA). They were fed with the same neuron chow and proliferation of non-neuronal cells was arrested by the addition of 1 µM cytosine arabinoside (Sigma-Aldrich, St. Louis, MO) at DIV 5. Cortical neurons were treated with 1 µM tetrodotoxin (TTX, Alomone Labs, Jerusalem, Israel) for 12 h, media was washed-out and bic/4AP were applied for 6 h at 21 DIV.

### HEK293T Cells Culture

HEK293T cells were cultured in DMEM media supported by fetal bovine serum.

### Experimental Animals

Neurons for primary cell cultures and slices for electrophysiology experiments were prepared from brains of Sprague Dawley (Janvier, France) or Wistar rats (Animal facilities of the Leibniz Institute of Neurobiology, Magdeburg, Germany). Male C57BL/6J (10-13 weeks old) were used for behavioral experiments (Charles River, / (Animal facilities of the Leibniz Institute of Neurobiology, Magdeburg, Germany). GluRε1 (GluN2A) knockout (KO) mice were obtained from RIKEN Japan (RBRC01813). Animals were housed in groups of up to 5 in individually ventilated cages (IVCs, Green line system, Tecniplast) under controlled environmental conditions (22 °C +/- 2 °C, 55% +/- 10% humidity, 12h light/dark cycle, with lights on at 06:00). Food and water were available ad libitum. All procedures and animal care were consented and performed under established standards of the German federal state of Sachsen-Anhalt, Germany in agreement with the European Communities Council Directive (2010/63/EU) and approved by the local authorities of Sachsen-Anhalt / Germany / Regierungspräsidium Halle Sachsen-Anhalt/Germany.

### Methylation analysis

DNA from the CA1 hippocampal tissue was extracted using Chargeswitch DNA extraction kit from Invitrogen (Carlsbad, CA, USA) according to instructor’s manual. Methylation-sensitive restriction enzyme-dependent methylation analysis was performed using OneStep q-Methyl Kit (Zymo, Irvine, CA, USA) following the instructions given in the manual using 20ng DNA for each sample. For methylated DNA immunoprecipitation (MeDIP) experiments the extracted DNA was then subjected to fragmentation by sonicator (Picoruptor, Diagenode, Seraing, Belgium) with 30 seconds on and 90 seconds off cycles using 1.5 ml Bioruptor microtubes. The efficiency of the sonication was confirmed by running the fragmented DNA samples on agarose gels and confirmed the accumulation of fragmented DNA was mainly at a range of 200-600bp. Afterwards MeDIP experiments were carried using the MagMeDIP (Diagenode, Seraing, Belgium) kit following the user guide with ∼750ng fragmented DNA as starting material. The methylation levels were assessed by qPCR and evaluated using the given formula in user guide of both kits. The forward primer used OneStep qMethyl-Kit was 5’-TATGACAGCTCACGTCAAGG-3’ and reverse primer 5’-CCTTCAGTGAGAAGCTCCAT -3’, containing three methylation sensitive restriction enzyme sites. The forward primer used in MeDIP-qPCR study for BDNF promoter IV was 5’-GCATGCAATGCCCTGGAACGG-3’ and reverse 5’-GAGGGCTCCACGCTGCCTTG-3’, and forward primer for BDNF promoter I 5’-TACCTGGCGACAGGGAAATC-3’ and reverse primer 5’-GCGCCCTAGCACAAAAAGTT-3’.

### Quantitative real time-PCR

14 DIV hippocampal neurons were treated with 50 µM bicuculline methiodide and 2.5 mM 4AP with or without 5nM MLN4924 for 6h. DMSO is used as sham control. Following the treatment, neurons were immediately harvested in lysis buffer and total RNA was isolated (RNeasy plus mini kit, Qiagen, Valencia, CA, USA). 50 nanograms of RNA were reverse transcribed using random nonamers (Sigma-Aldrich, St. Louis, MO, USA) according to the manufacturer’s instructions (Sensiscript, Qiagen). BDNF exon IV and glyceraldehyde 3-phosphate dehydrogenase (GAPDH) mRNA (as a reference gene) were amplified using the iScript RT-PCR iQ SYBR Green Supermix (BIORAD, Hercules, CA, USA) in a real-time quantitative PCR (qPCR) detection system (LC480, Roche, Basel, Switzerland) using the following primers: *Bdnf* exon IV forward 5’-GCAGCTGCCTTGATGTTTAC-3’ and reverse 5’-CCGTGGACGTTTGCTTCTTTC-3’; *Dnmt3a1* forward 5’-CCCTCAGATCTGCTACCCAA-3’ and reverse 5’-TGCTCTGGAGGCTTCTGGTG-3’; *Gapdh* forward 5’-TGCTGAGTATGTCGTGGAG-3’ and reverse 5’-GTCTTCTGAGTGGCAGTGAT-3’. Each sample reaction was run in duplicates and Ct values of the reference genes from the samples were subjected to Grubbs’ outlier test (http://www.graphpad.com/quickcalcs/Grubbs1.cfm). The relative expression levels were analyzed using the 2-ΔΔCt method with the normalization relative to GAPDH.

### Immunocytochemistry

Following drug treatment, cells were fixed either in methanol or in 4% Paraformaldehyde (PFA) and immunostaining was performed with primary antibodies specific for DNMT3AN-terminus (generous gift from Prof. Tajima, Osaka, Japan), CUL4B (Proteintech, catalog #12916-1-AP), GFAP (Synaptic Systems, catalog #173011), HOMER1 (Synaptic Systems, catalog #160011), NEDD8 (Proteintech, catalog #16777-1-AP), MAP2 (Sigma-Aldrich, catalog #M4403), 5meCytosine (Calbiochem, catalog #MABE146, clone 33D3), Synaptophysin1 (Synaptic Systems, catalog #101004); secondary antibodies anti-rabbit/mouse-Alexa Fluor 488/568 linked (Molecular Probes Europe BV, Leiden, The Netherlands), anti-rabbit/mouse-Cy™5-conjugated (Dianova, Hamburg, Germany) and DAPI (Sigma-Aldrich, catalog #D9564). Live immunstaining was performed for the detection of the surface expression of 2A subunit of NMDARs, therefore, GluN2A antibody (Alomone Labs, #AGC-002) was applied in cell media for 20 minutes, then cells were fixed, and staining protocol was followed as mentioned above.

### Confocal laser scanning microscopy and image analysis

The SP5 CLSM system (Leica-Microsystems, Mannheim, Germany) equipped with Diode (405nm) and Argon (488, 561, 633nm) lasers lines was used for quantitative immunocytochemistry. Z-stack images of neurons were taken using the 63x oil-immersion objective (Leica, Mannheim, Germany). Confocal images of triple-stained neurons were taken with Plan Apo 63x oil NA 1.4 objective lenses. All images were acquired sequentially to avoid crosstalk between channels. The acquisition parameters were kept the same for all scans. Regions of interest (ROI) were drawn around nuclei, as delineated by DAPI staining. These ROI were then applied to a corresponding image of antibody staining from which the mean average intensity was collected to determine nuclear immunoreactivity levels using Image J software (NIH, Rasband, W.S., ImageJ, U. S. National Institutes of Health, Bethesda, Maryland, USA, http://imagej.nih.gov/ij/, 1997-2014).

Total number of dendritic spines per 20 µm secondary dendrites of the primary hippocampal neurons, which received 5 nM or 1 µM MLN4924 treatment at basal conditions or with synaptic stimulation using bic/4AP were counted based on the Homer1 staining. Total synapse number per 20 µm secondary dendrites of the primary hippocampal neurons following Cul4B knockdown were counted based on the colacalization of Homer1 and Synaptophysin1. Surface expression of GluN2A was evaluated in primary hippocampal neurons following the knockdown of Cul4B. Total number of of GluN2A staining per 20 µm secondary dendrites of primary hippocampal neurons were counted.

### Fluorescent microscopy and image analysis

The number of dendrites crossing each circle was counted manually. Studies of GluN2A knockdown on hippocampal primary hippocampal neurons were also carried out using Zeiss Axio Imager A2 fluorescent microscope (Zeiss, Jena, Germany) with Cool Snap EZ camera (Visitron System) and MetaMorph Imaging software (MDS Analytical technologies). Up to 3 coverslips were treated individually and processed per group. For each coverslip among the different groups, the same exposure time and intensity were taken. Upon background subtraction using Fiji software, images of fluorescent positive puncta were measured along secondary dendrites, right after the brunching pint. The synaptic immunofluorescence intensities were assessed in a region of 400 nm x 400 nm square set by the mask in the channel for Homer1, used as synaptic marker. The mask was created semiautomatically using Openview software. The intensities of the puncta positive for Glun2A were measured at the colocalization spots with Homer1; values were normalized to control mean and plotted.

### Intrahippocampal injections

Mice were anesthetized with 5% isofluorane in O2/N2O mixture. Mice were placed in a stereotaxic frame (World Precision Instruments) and anesthesia was maintained with 1.5% isofluorane using gas anesthesia system (Rothacher Medical GmbH., Switzerland). After craniotomy, 10 µl NanoFil microsyringes (World Precision Instruments) containing 33G injection needles were lowered into the dorsal CA1 area under stereotactic guidance with the coordinates anterioposterior (AP) −2.0 mm, mediolateral (ML) ±1.5 mm from Bregma and dorsoventral (DV) −0.14 mm from brain surface. Each animal received 1.5 µl/hemisphere bilateral infusion of drug (MLN4924) while (saline with respective volume of DMSO) as sham control, at an infusion rate of 0.5 µl/min.

### Overexpression and knockdown experiments

The overexpression experiments in HEK-293T cells were performed using the following constructs: C2-eGFP-hDNMT3A (Clontech, subcloned from pcDNA3/Myc-DNMT3A1, which was a gift from Arthur Riggs, Addgene plasmid #35521 [4], C2-eGFP-hDNMT3A2 (Clontech, subcloned from pcDNA3/Myc-DNMT3A2, which was a gift from Arthur Riggs, Addgene plasmid #3694, pK-MYC-C3 (gift from V. Horejsi Lab, Prague, Czech Republic), pcDNA3-myc3-CUL4B (gift from Yue Xiong, Addgene plasmid #19922, pcDNA3-myc3-NEDD8 (gift from Yue Xiong, Addgene plasmid #19943, pcDNA3-HA-NEDD8 (gift from Edward Yeh, Addgene plasmid #18711, CUL1, CUL2, CUL3, CUL4A, CUL4B, CUL5 and CUL7 in pcDNA3-myc3 backbone were a gift from Yue Xiong (Addgene plasmid #19896, #19892, #19893, #19951, #19922, #19895, #20695 respectively). The shRNA constructs to knockdown human DNMT3A1/3A2 and scrambled controls were hDNMT3A shRNA pSMP-DNMT3A1 and pSMP-Luc (a gift from George Daley, Addgene plasmid #36380, Addgene plasmid #36394, respectively. shRNA targeting, ratDNMT3A and scrambled control were cloned into pZ-off vector with the following sequences: CCCAAGGTCAAGGAGATCA and GCTTCGCGCCGTAGTCTTA, respectively. shRNA targeting rat-Nedd8, human-Nedd8 and scrambled control were cloned into pSuper vector with the following sequences: GCGGCTCATCTACAGTGGCAA, GAGGCTCATCTACAGTGGCAA, CTTCGCGCCGTAGTCTTA, respectively.

### Acute hippocampal Slice Preparation and Electrophysiology

Hippocampi from 8 weeks old male Wistar rats (House strain, LIN, Magdeburg) were cut using a vibratome (LeicaVT1000S, Nussloch, Germany) into 350 µm thick slices. The hippocampal slices were incubated for 2h in carbogenated (95% O2 ∼ 5%CO2) artificial cerebrospinal fluid (ACSF, containing in mM: 110 NaCl, 2.5 KCl, 2.5 CaCl2·2H2O, 1.5 MgSO4·7H2O, 1.24 KH2PO4, 10 glucose, 27.4 NaHCO3, pH 7.3) at room temperature. Then slices were transferred into a recording chamber (at 31±1 °C). Field excitatory postsynaptic potentials (fEPSPs) were evoked by stimulation of CA1 Schaffer-collateral fibers with metal electrodes. fESPSs were recorded with ACSF filled glass capillary microelectrodes (3-5 MΩ) and amplified by an Extracellular Amplifier (EXT-02B, npi, Germany) and digitized at a sample frequency of 5 kHz by Digidata 1401plus AD/DA converter (CED, England). Stimulation strength was adjusted to 30% ∼ 40% of the maximum fEPSP-slope values. Single stimuli were applied every 30 s (at 0.0333 Hz) and were averaged every 3 min. After 30min stable baseline recording, MLN4924 was applied into bath. 30 min later, late-long term potentiation (L-LTP) was induced by tetanization consisting of three 1 s stimulus trains at 100 Hz with a 6 min inter-train interval where the width of the single stimulus was 0.1ms. MLN4924 was dissolved in DMSO and diluted in ACSF at final concentration (5 μM, or 50 nM) applied 30 min before the first tetanus. Data are represented as mean ± SEM. The preparation of acute hippocampal slices from GluN2A KO mice (17-22weeks) was the same as Wistar rats (8 weeks), the method of electrophysiology test as well, but the LTP recording time was 3h on N2A mice (shorter than 6h on Wistar rats).

### Tissue collection and analysis

After the LTP recordings, only the potentiated CA1 region of the Hippocampus was collected from the acute slices. The tissue was either subjected to RNA or DNA extraction for transcription or promoter methylation analysis, respectively, or for total protein extraction for western blotting. For the experiments investigating learning-dependent DNMT3A1 degradation, mice were sacrificed 3 or 6 hours after training and CA1 region of the hippocampus was dissected. Bilateral intra-hippocampal MLN4924 (7.5 pmol/site) or saline infusions were performed immediately after training.

### Immunoprecipitation experiments

Endogenous IP experiments were performed from cultured rat cortical neurons. 50 µl dynabeads Protein G (ThermoFischer Scientific, Waltham, MA, USA) were blocked using albumin from chicken egg white (Sigma-Aldrich, St. Louis, MO, USA) while rocking for 30 min. Then, the dynabeads protein G were incubated with 2-3 µg of rabbit polyclonal CUL4B antibody (Proteintech, catalog #20882-1-AP) while gently rotating for 2h at 4°C. Nuclear protein extracts were then incubated with antibody-bound-protein G dynabeads overnight at 4 °C. The next day, following several washes with ice-cold TBS-T, samples were eluted using 4x Laemmli sample buffer. Heterologous co-immunoprecipitation (het-coIP) experiments were performed in HEK-293T cells. Transfection of HEK-293T cells was performed with constructs pcDNA3-myc3-CUL4B, C2-eGFP-hDNMT3A, and pK-MYC-C3. 3h prior to harvesting the cells either the proteasome inhibitor MG132 (30 µM) or DMSO as sham control was applied on the cells. 24h after transfection cells were harvested. In the het-coIP experiment in which the effect of neddylation was studied, transfections were performed using the constructs pcDNA3-myc3-CUL4B, C2-eGFP-hDNMT3A, pK-MYC-C3, and myc3-NEDD8. 24h after transfection HEK-T cells were treated with NEDD8-activating enzyme inhibitor MLN4924 (94nM) or DMSO as sham control for 24 h. MG132 application was done as mentioned above. Cell lysate extracts were incubated with the anti-GFP microbeads (Miltenyi Biotec GmbH, Gladbach, Germany) for 1h. Eluted samples were run on SDS-PAGE.

### Western blot

Total homogenates were prepared from mouse and rat brain or cultured cortical neurons by performing the lysis in 10mM Tris/HCl pH 7.5, 0.5% TritonX, and protease inhibitor cocktail (Roche, catalog #04693116001) containing TBS. 4x Laemmli buffer was added to a final dilution of 1.5x following 10 min-incubation at 95°C, samples were ready for protein analysis. HEK-293T cells were harvested in TBS, which contains protease inhibitor cocktail. Cells were lysed in 1% Triton-X containing lysis buffer; centrifuged for 30 min at 21000 rpm, the supernatant fraction was collected. Protein estimation was performed either by amidoblack or BCA assay (Thermo Scientific). Western blots were then performed using 4-20% gradient polyacrylamide gels. The following antibodies were used in this study: DNMT3AN (1:2000) and DNMT3A-mid (1:2000), CUL4B (1:1000, Proteintech, catalog #12916-1-AP), NEDD8 (1:1000, Proteintech, catalog #16777-1-AP), FK1 (1:1000, Enzo, catalog #BML-PW8805), β-Actin (1:2000, Sigma-Aldrich, catalog #A5441), myc-tag (1:1000, Cell Signaling, catalog #2276), and horseradish peroxidase-coupled goat anti mouse/rabbit IgG-HRP linked secondary antibodies (1:20000, Jackson Immunoresearch Laboratories, catalog #115035003 and #111035003, respectively). Quantification of immunoblots was done with ImageJ software (NIH, Maryland, USA). Integrated density values were evaluated for the analysis. When necessary, experimental data from individual experiments performed at different time points were normalized to respective controls and pooled together with other data set.

### Object Location Memory

Object location memory was performed in a square arena (50 × 50 × 50 cm) under mild light conditions, according to Heyward and colleagues (2012), with modifications. Briefly, the task consisted of a habituation session, training and test. During habituation, animals were allowed to explore the empty arena for 20 minutes. Twenty-four hours later, training session took place, where animals were free to explore a pair of similar objects (made of plastic mounting bricks), placed in the arena, for 20 minutes. Test session was performed 6 hours after training, where one of the objects was was placed in a new position, and again, animals were free to explore the two objects for 20 minutes. All 3 sessions were video-recorded and behavior was analyzed offline using ANY-maze software (ANY-mazeTM Video Tracking System, version 4.50, Stoelting Co. Wood Dale, USA). Exploration was recorded only when the animal touched or reached the object with the nose at a distance of less than 2 cm. The time mice spent exploring the objects was recorded, and discrimination index was then calculated, taking into account the difference of time spent exploring the new and familiar position ([(Tnovel – Tfamiliar)/(Tnovel + Tfamiliar)] × 100). An experimenter blind to treatment conducted experiment and data analysis. Chambers and objects were thoroughly cleaned with 10% ethanol before and after each animal was test.

### Statistical Analysis

The data were analyzed by one-way ANOVA and unpaired one/two-tailed Student’s t-test. Two-way ANOVA followed by post-hoc Bonferroni test was employed to compare means from multiple groups. Quantitative real-time PCR data were subjected to Grubbs’ outlier test and analyzed by either unpaired two-tailed Student’s t-test or two-way ANOVA, which was followed by Bonferroni post hoc test, where applicable. The Mann Whitney U-test was used to compare the Averaged field potentials (300-360 min) between two groups of differentially treated slices. Otherwise stated, error bars present S.E.M. Statistical analyses were performed in GraphPad (GraphPad Software, Inc., La Jolla, USA).

## Acknowledgments

We would like to thank Monika Marunde, Corinna Borutzki, and Stefanie Hochmuth for excellent technical assistance. We are also grateful to Dr. Anna Karpova for help in preparation of the Figures. We would like to thank Masayoshi Mishina from the University of Tokyo (Graduate School of Medicine) for the generous gift of GluRε1 (GluN2A) KO mice.

Supported by grants from the Deutsche Forschungsgemeinschaft (DFG Kr1879 / 5-1 / 6-1 / SFB 779 TPB8), BMBF ‘Energi’ FKZ: 01GQ1421B, The EU Joint Programme – Neurodegenerative Disease Research (JPND) project STAD and Leibniz Foundation to MRK; Deutsche Forschungsgemeinschaft SFB 779 TPB5 to OS. Grants-in-Aid for Scientific Research B from the Japan Society for the Promotion of Science to ST; Deutsche Forschungsgemeinschaft EXC 257/2 to FY. GMG was supported by a CAPES-Alexander von Humboldt Research Fellowship (1756/14-1)

## Author contributions

GB designed, performed experiments, analyzed data and wrote the manuscript. PY and ADR performed experiments and analyzed data. GMG, SAR, OS designed, performed and analyzed data from behavioral experiments. ST provided and characterized the DNMT3AN and DNMT3A-mid antibodies. FY designed and supervised DNA methylation analysis experiments. MRK designed and supervised the study, and wrote the manuscript.

## Expanded View Figures

**Figure EV1.**
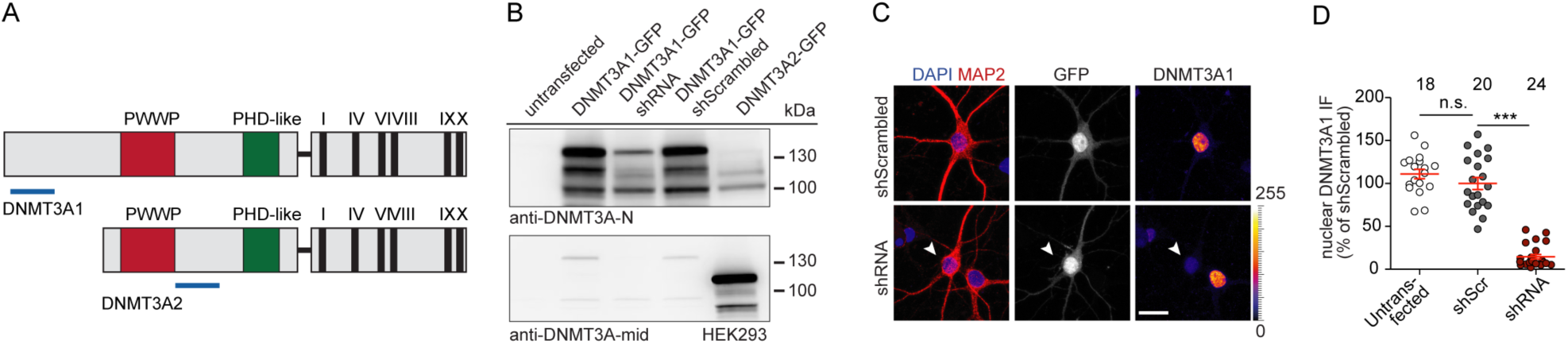
Verification of the specificity of the DNMT3A isoform1 antibody. **A** The domain structure of the two *de novo* DNA methyltransferase DNMT3A1 and DNMT3A2 is depicted. The epitopes of the DNMT3AN and DNMT3A-mid antibody are indicated with blue bars. **B** Heterologous expression of DNMT3A1 and DNMT3A2 and shRNA-based mRNA knockdown in HEK-T cells was detected with either the Dnmt3aN specific antibody (upper panel) or the DNMT3A-mid antibody recognizing both isoforms (lower panel). **C** and **D** Representative images and quantification depicts the nuclear DNMT3A1 isoform knockdown in 16 DIV primary hippocampal neurons. Unpaired two-tailed Student’s t-test, n.s. not significant, *** p<0.0001. Scale bar is 20 µm.

**Figure EV2.**
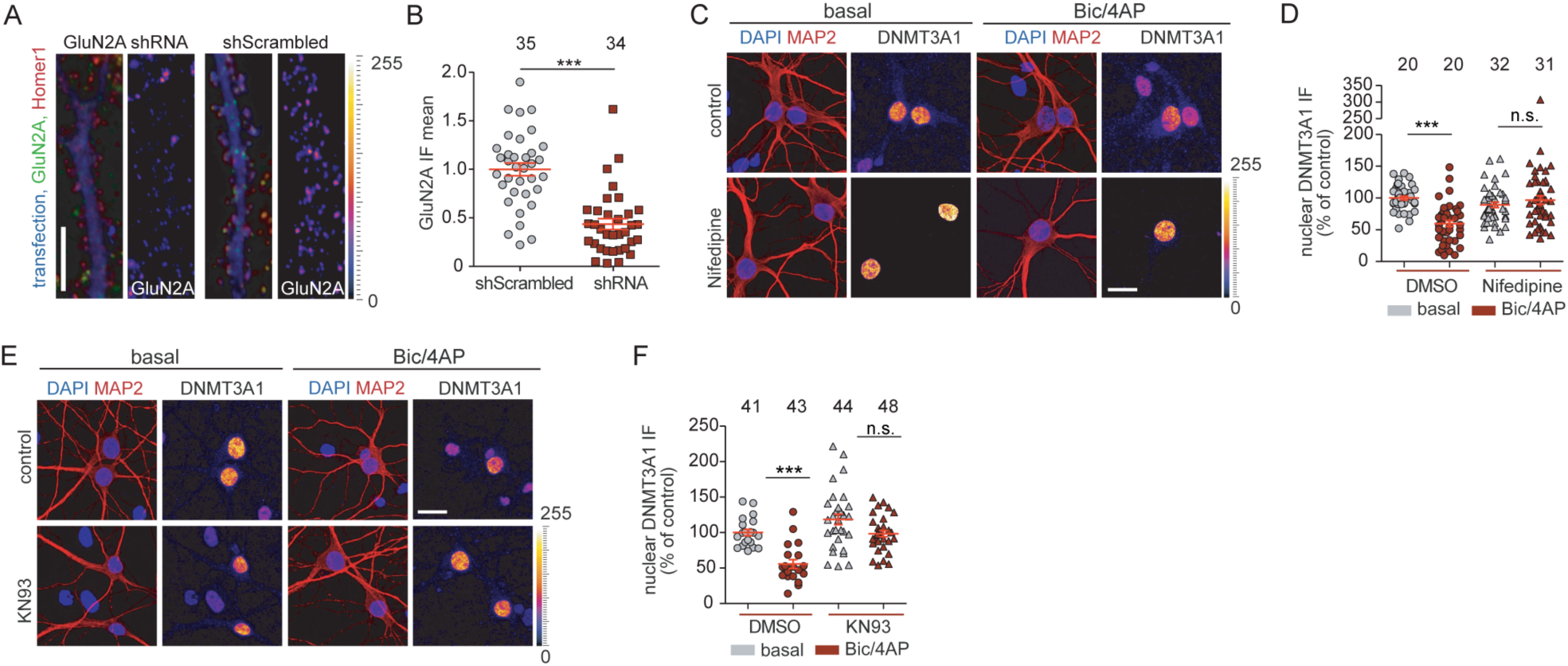
KN93 and Nifedipine attenuates DNMT3A1 degradation upon neuronal stimulation. **A** and **B** Representative immunofluorescence images of 11 DIV hippocampal primary neurons transfected with either shScrambled or GluN2A shRNA, fixed after 4 days after transfection at 15 DIV. shRNA knockdown confirms the significant less surface expression of GluN2A. Scale bar is 10 µm. Unpaired two-tailed Student’s t-test. **C** and **D** The L-type calcium channel blocker Nifedipine (10 µM) attenuated the degradation of DNMT3A1 when 14-15 DIV hippocampal primary neurons were stimulated by bic/4AP for 6h. Two-way ANOVA followed by Bonferroni’s post hoc test. Scale bar is 20 µm. **E** and **F** The CamKII inhibitor KN93 (5 µM) abolished the degradation of DNMT3A1 when 14-15 DIV hippocampal primary neurons were stimulated by bic/4AP for 6h. Scale bar is 20 µm. Two-way ANOVA followed by Bonferroni’s post hoc test. ***P<0.001, n.s. not significant.

**Figure EV3.**
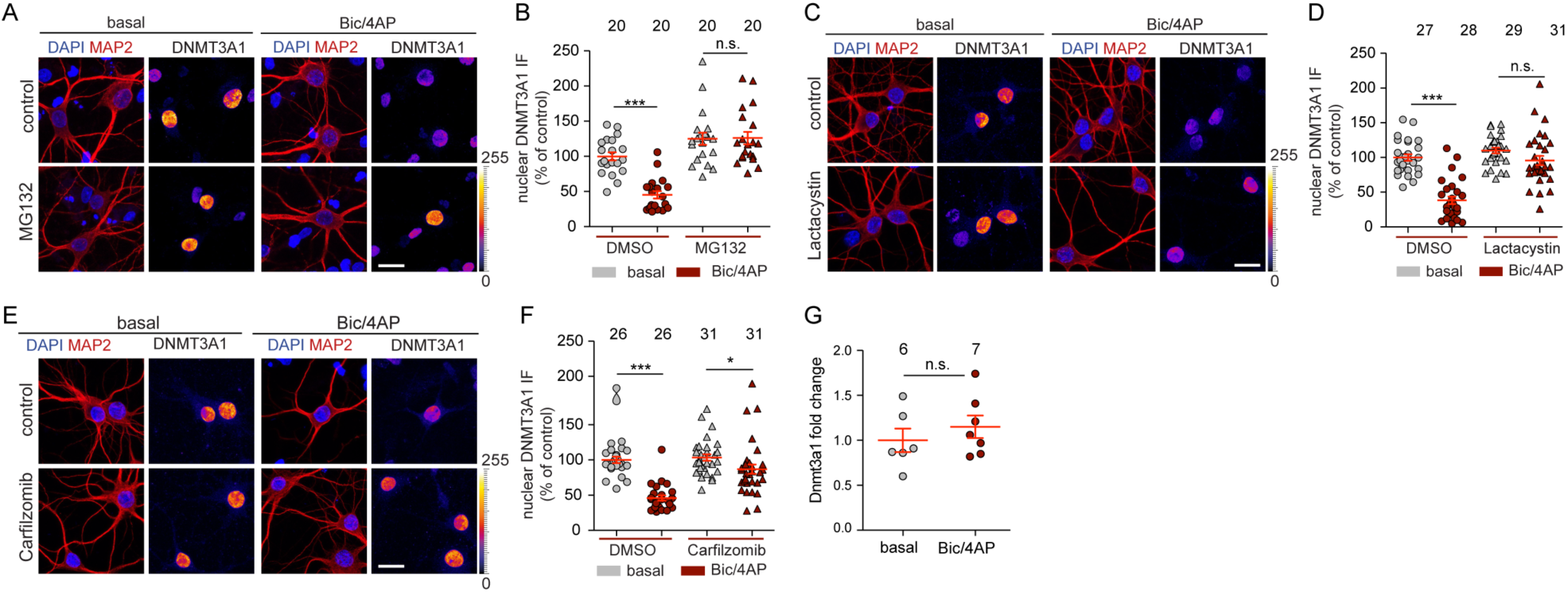
Proteasomal degradation of DNMT3A1 following synaptic stimulation. **A** and **B** Inhibition of the proteasome with 30 µM MG132, **C** and **D** 15 µM Lactacystin and **E** and **F** 100 nM Carfilzomib prevented degradation of DNMT3A1 upon neuronal stimulation. Scale bars are 20 µm. Two-way ANOVA followed by Bonferroni’s post hoc test. ***P<0.001, n.s. not significant.

**Figure EV4.**
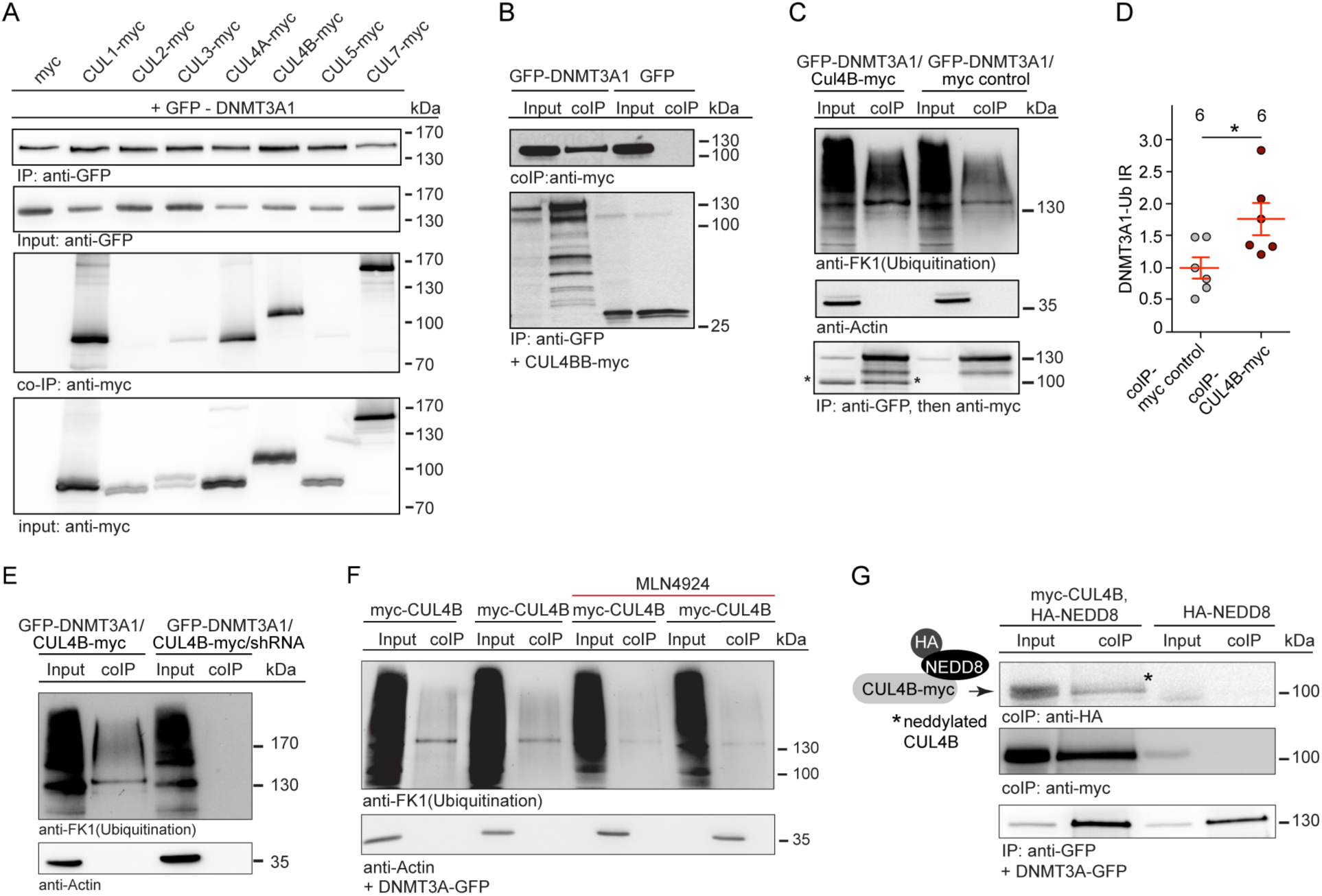
Neddylation of the Cul4B enhances DNMT3A1 degradation. **A** Heterologous expression of GFP-DNMT3A1 and empty myc-vector or human myc-Cullins (CUL1, CUL2, CUL3, CUL4A, CUL4B, CUL5, CUL7) in HEK293-T cells. DNMT3A1 was immunoprecipitated from total cell extracts using anti-GFP antibodies coupled to microbeads. Cullin proteins were detected in the immunoprecipitate with a myc-antibody. Same volume of IP and input per sample were loaded. **B** HEK-T cells were transfected with expression vectors for GFP-DNMT3A1 and myc-CUL4B or myc-vector. DNMT3A1 was immunoprecipitated from total cell extracts using anti-GFP antibodies coupled to microbeads. CUL4B was detected in the immunoprecipitate with a myc-antibody. **C** and **D** Poly-ubiquitination of DNMT3A1 in the immunoprecipitate is enhanced when DNMT3A1 was co-expressed with CUL4B. Unpaired two-tailed Student’s t-test, *p<0.05. **E** shRNA knockdown of CUL4B reduces polyubiquitination of DNMT3A1. **F** and **G** HEK-T cells were transfected with expression vectors for GFP-DNMT3A1, myc-CUL4B (**F**) and also with active HA-Nedd8 (**G**). Total cell extracts were immunoprecitated using anti-GFP antibodies. (**F**) Less DNMT3A1 ubiquitination was observed when the NAE inhibitor MLN4924 (94nM) was applied for 24h. (**G**) Neddylated CUL4B was detected with anti-HA antibody in the DNMT3A1 elute.

**Figure EV5.**
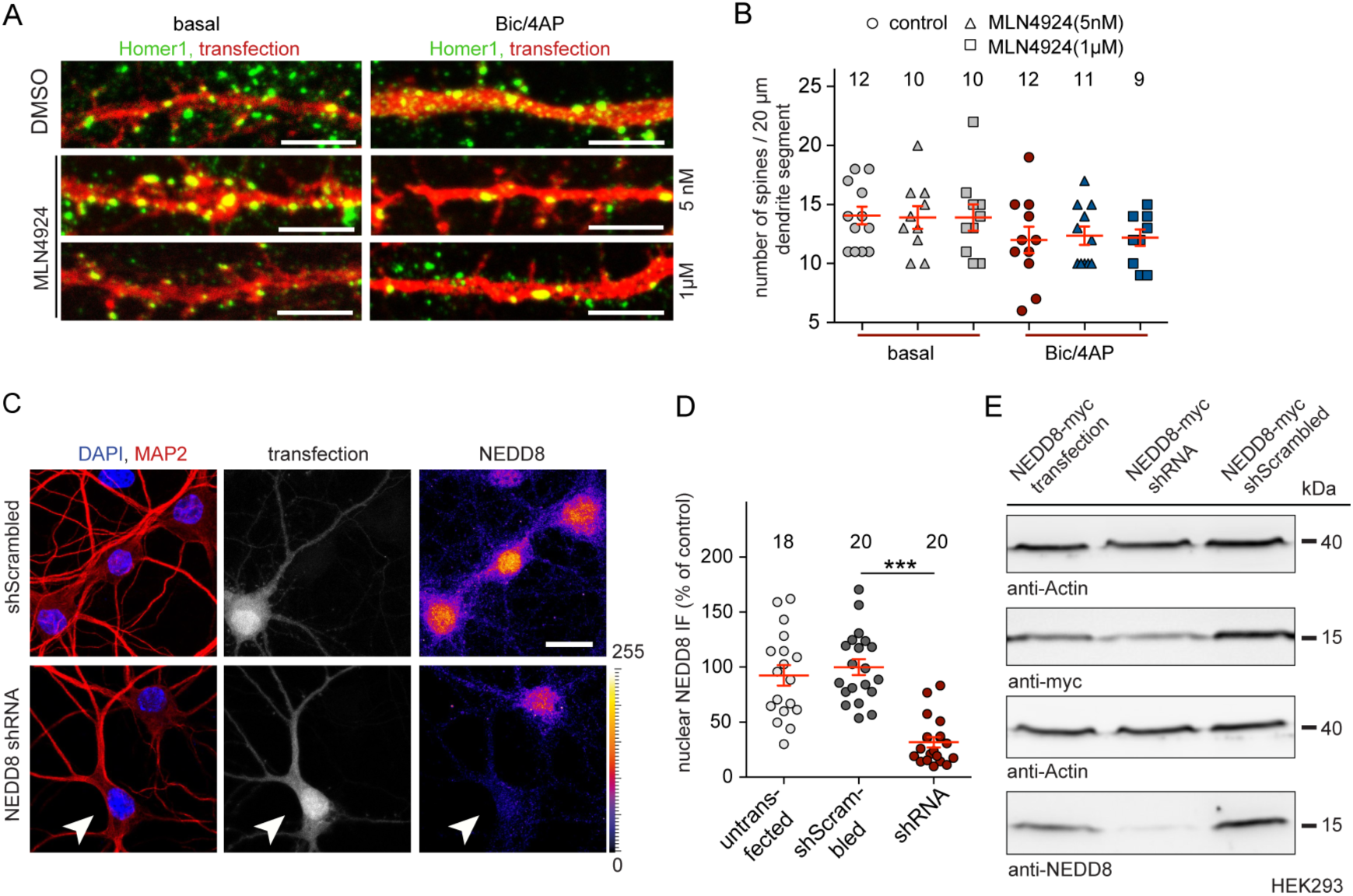
Inhibition of neddylation does not alter total spine number. **A** and **B** DIV 14 hippocampal neurons were transfected with RFP expressed under control of an actin promoter and cells were fixed two days after transfection. Homer1 positive dendritic spine numbers were counted. There is not any significant alteration in the total spine number following MLN4924 treatments following the synaptic stimulation with bic/4AP for 6 h. Scale bars are 5 µm. **C** and **D** Representative images and quantification display the endogenous rat-Nedd8 knockdown generated in 16 DIV primary hippocampal neurons. Unpaired two-tailed Student’s t-test, *** p<0.0001. Scale bar in C is 20 µm. **E** Heterologous expression of human-NEDD8 and shRNA mRNA knockdown and shScrambled control in HEK-T cells was detected with either the anti-myc antibody (upper panel) or the anti-NEDD8 antibody (lower panel). The knockdown of Nedd8 was detected by anti-myc and as well as anti-NEDD8 antibodies.

**Figure EV6.**
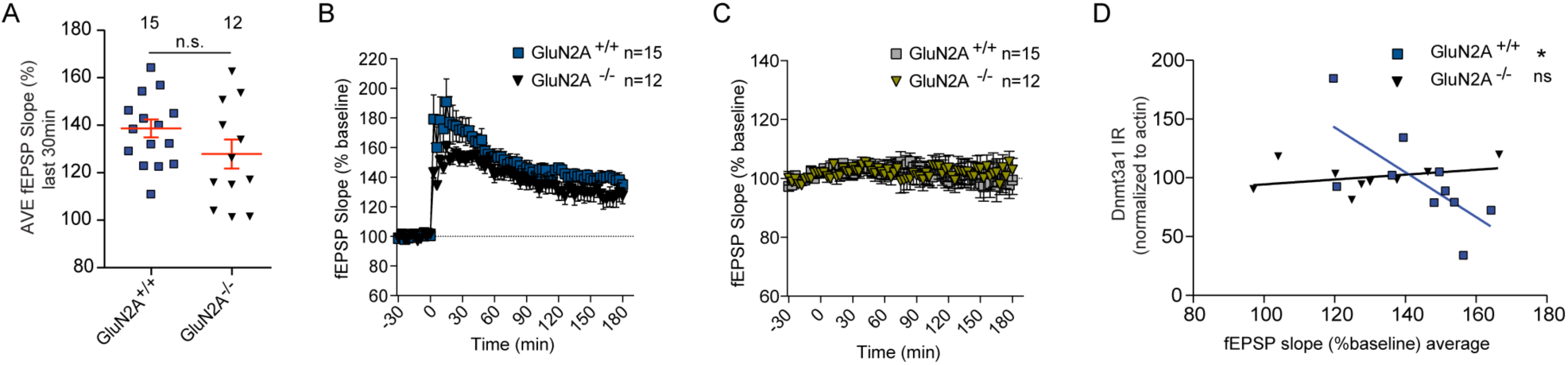
Recovered LTP in GluN2A knock-out mice. **A** Averaged fEPSP slopes of the last 30 min LTP showed no significant difference between GluN2A+/+ and GluN2A−/− mice. The Mann Whitney Test was used to compare the Averaged field potentials (150-180min) between two groups. **B** LTP induced by HFS (3 trains high frequency stimulus @100Hz) in GluN2A−/− mice (n=12 slices from 6 animals) was similar to GluN2A+/+ mice (n=15 slices from 6 animals). **C** Baseline was determined by the second input on the same slice which was induced LTP by the first input. **D** Scatterplot and correlation analysis between the normalized DNMT3A1 immunoblotting mean values and the averaged fEPSP slope of the whole LTP recording depicts a negative correlation between the strength of the LTP and the synaptic stimulation-induced DNMT3A1 down-regulation in GluN2A +/+ mice while no significant correlation was observed in slices of GluN2A −/− mice. Unpaired two-tailed Student’s t-test, *P<0.05, ns not significant.

**Figure EV7.**
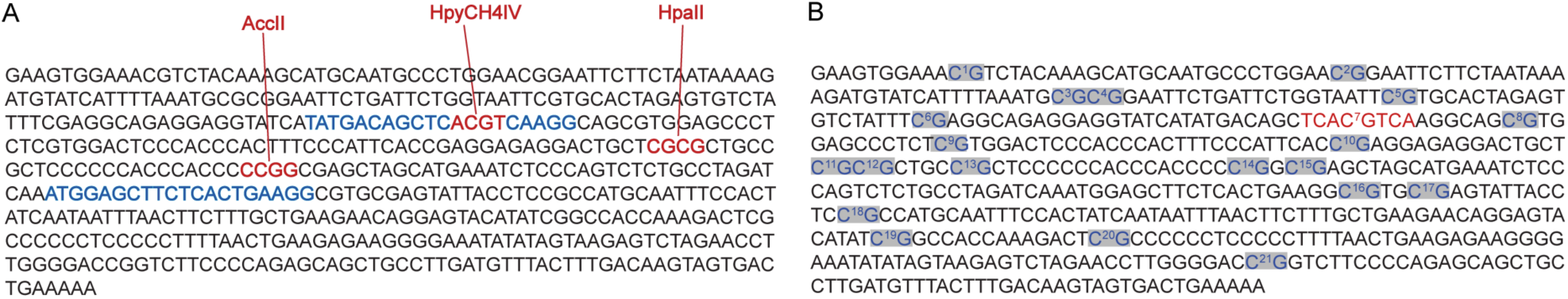
The primers used and CGs covered in *Bdnf* promoter methylation analysis. **A** *Rattus norvegicus* Bdnf IV promoter sequence (chr3:100,786,676-100,787,200 on the UCSC genome browser) shows: Primers, highlighted in blue, used in the study spanning a sequence, which contains three methylation-sensitive restriction sites, depicted in red. *Rattus norvegicus* Bdnf IV promoter sequence (chr3:100,786,676-100,787,200 on the UCSC genome browser) shows: CG dinucleotides highlighted in grey and CRE binding site in red. **B** *Rattus norvegicus* Bdnf IV promoter sequence (chr3:100,786,676-100,787,200 on the UCSC genome browser) shows: CG dinucleotides highlighted in grey and CRE binding site in red.

